# Neuromechanical Coupling is Reflected in the Spatial Organization of the Spinal Motoneuron Pools

**DOI:** 10.1101/2022.07.09.499432

**Authors:** Rachel I. Taitano, Sergiy Yakovenko, Valeriya Gritsenko

## Abstract

Recent advances in neuroprosthetics have shown that activating the surviving spinal circuitry below the level of spinal damage can produce complex movements and restore function. This is because the spinal cord circuitry plays an important role in the control of complex movements by integrating descending execution-related signals with feedback sensory signals. To generate efficient movements through muscle contractions, the nervous system embeds the anatomical and dynamical properties of the controlled body parts, a concept termed neuromechanical tuning. The motoneurons innervating the skeletal muscles of the body form complex spatial structures that span multiple segments. The goal of this study is to determine if these structures can support neuromechanical tuning. We performed a comparative analysis of the spatial organization of the motoneuron pools and the functional organization of the muscles they innervate. We developed a three-dimensional model of the spatial organization of motoneuron pools innervating upper limb muscles in the macaque and quantified their relative distances. We then utilized a musculoskeletal model of the macaque upper extremity to characterize the functional relationship between the muscles innervated by the motoneuron pools. We found that the distances between motoneuron pools mimicked the agonistic or antagonistic actions of muscles they innervate. This shows that the spatial organization of the motoneuron pools embeds the functional musculoskeletal anatomy evolved to support the large repertoire of upper limb movements in primates. This spinal anatomy may play an important role in the neuromechanical tuning and simplify the sensorimotor control of the arm.

## Introduction

Epidural electrical stimulation is the most promising technique to restore functional movement after spinal cord injury ^1–8^. Models of the spinal cord anatomy play a crucial part in surgical planning and tuning of the neuroprosthesis for the needs of the individual ^7^. Here we developed a model of the anatomical arrangement of spinal motoneuron (MN) pools of the upper limb and related it to the function of the muscles that MN pools innervate.

The spinal cord drives the musculoskeletal system by rapidly integrating ongoing execution-related (efferent) signals with proprioceptive (afferent) signals to enable motor function in presence of uncertain environmental conditions. Preserving and taking advantage of proprioceptive signals is crucial for the optimal augmentation of voluntary movement after spinal cord injury using neuroprostheses ^3^. The afferent signals are somatotopically organized carrying information from specific body parts to separate spinal cord segments ^9,10^. We also know that the primary motor cortex, one of the main generators of the efferent signals for the motor control of the upper limb, is somatotopically organized ^11–14^. This organization effectively embeds the anatomical and dynamical properties of the controlled body parts in the neural substrate, a concept referred to as internal models or, specific to the embedding of the musculoskeletal system, neuromechanical tuning ^15–18^. The neuromechanical tuning serves to simplify the processing of the proprioceptive signals for the generation of efficient efferent commands. The question remains as to how much the spinal cord contributes to the neuromechanical tuning.

Answering this question will help develop new methods to tune the neuroprosthetics for maximal effectiveness.

Comparative analysis between the MN pool anatomy and the function of muscles they innervate can help understand how the location of MN pools help integrate the afferent and efferent neural commands. The general understanding of the anatomical organization of MN pools innervating the human leg and some arm muscles comes from postmortem studies ^19–21^. There is also detailed anatomical data of the distribution of MN pools innervating muscles of the lower limbs in cats obtained with retrograde staining ^22^, which has been comprehensively analyzed ^23^. We also know that the rostrocaudal distributions of these MN pools are similar to those in humans ^21,24^.

However, the detailed anatomical data about the distribution of MN pools innervating the upper limb muscles in humans is sparse and lacks identification of their targets ^19^. In this study we used the detailed anatomical organization of MN pools innervating the upper limb muscles of macaques obtained with retrograde staining ^25^ to create a spatial model of the MN distributions in the spinal cord ventral horn. We then took advantage of the detailed information about the organization of the musculoskeletal system developed using advanced imaging and modeling techniques ^16,26,27^. Our goal is to determine if there is an anatomical substrate in the spinal cord that can support neuromechanical tuning. We will achieve this goal through the comparative analysis of the spatial organization of the MN pools and the functional organization of the muscles they innervate.

## Materials & Methods

All data processing and analyses were done in Matlab (Mathworks, Natick, MA).

### Model of the Spatial Organization of Motoneuron Pools

Locations of motoneuronal cell bodies innervating macaque forelimb muscles were acquired from a study utilizing retrograde transport of horseradish peroxidase in 10 *M. mulatta* and 4 *M. fascicularis* monkeys ^25^. Images of spinal cord sections from the rostral and caudal parts of the 5^th^ cervical segment (C5) through the 1^st^ thoracic segment (T1) were scanned, digitized using Adobe Illustrator (Adobe Inc) and imported into Matlab. Metadata from these images representing cell body locations in the three orthogonal planes (mediolateral, dorsoventral, and rostrocaudal axes) and the segmental outlines of white and gray matter were used to generate a three-dimensional (3D) model, similar to our previous work (Yakovenko et al., 2002). One representative set of transverse grey matter outlines (*M. fascicularis* subject 79-1 from Jenny and Inukai ^25^) was selected as the standard in our model and used for the spatial normalization of both outlines and MN coordinates along the dorsoventral and mediolateral axes. The published data for the anatomical size of spinal segments in monkeys ^28^ was used to scale the rostrocaudal coordinates of labeled MNs.

The digitization of MNs from published bitmap images of spinal cord section was challenged by image resolution, color representation, and the superposition of cells in the plane of each slice and the resolution. To mitigate this, additional data from the published histograms of the quantity of MNs in each section was utilized ^25^. The MN locations obtained from transverse sections were resampled to match the numbers reported in the histograms for each spinal cord section.

Briefly, if the dataset extracted from transverse sections included higher number of MNs than reported in the histogram for the corresponding pool in a given section due to mislabeling of irregular shapes in pixelated images, then MN indices for the given MN pool and section were randomly selected for removal. Conversely, the reduction in the number of digitized MNs was also possible due to the potential omission of some images of the transverse sections in the manuscript figures. This was compensated by adding missing MNs in accordance with the spatial distribution of the given MN pool and section. The mean position and standard deviation within transverse slices and the range across spinal segments were held constant for each MN pool.

Symmetry was assumed between left and right sides of the spinal cord when combing data from different animals to generate a bilateral model (Fig. 1).

**Figure 1.**
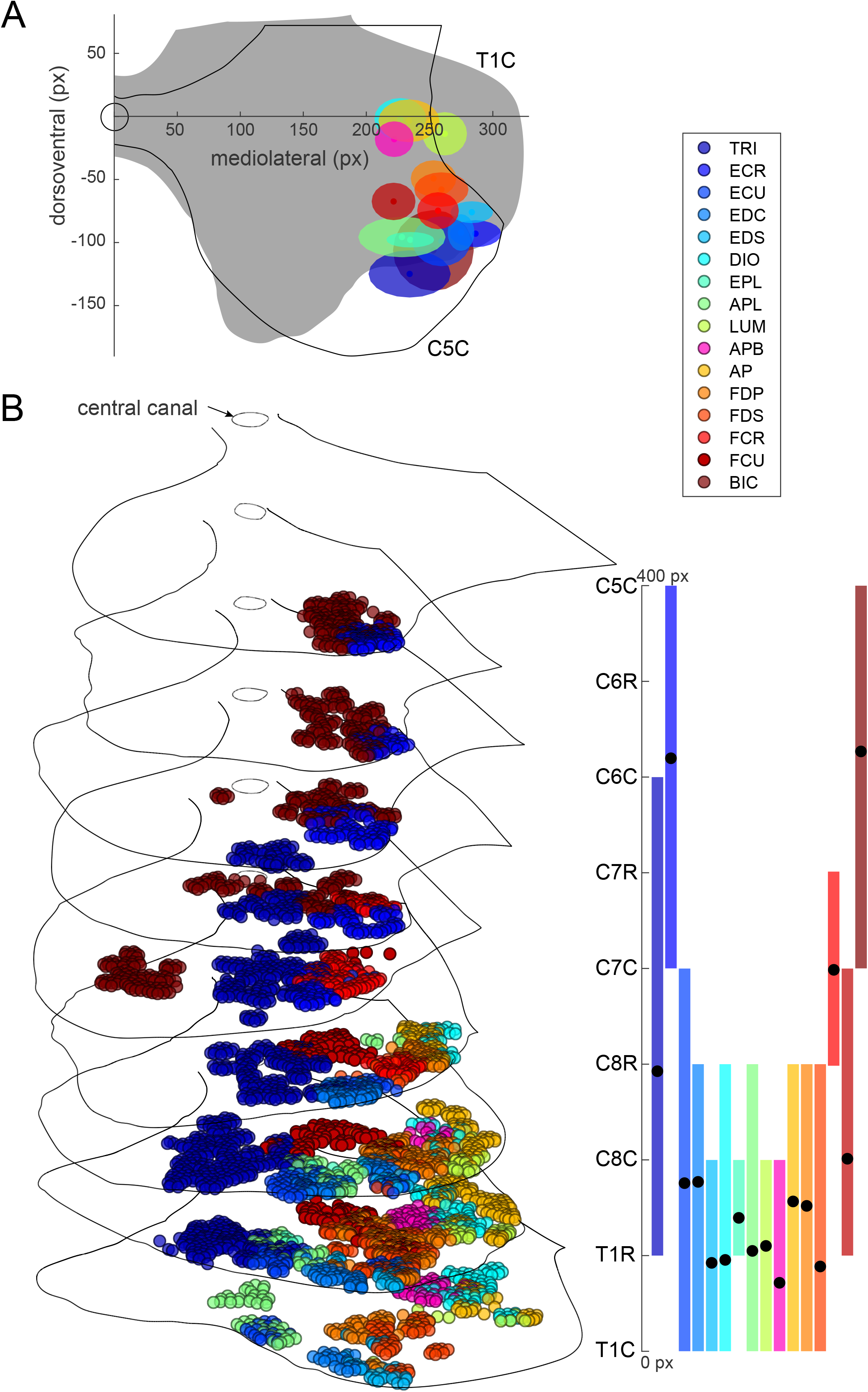
Model of motoneuron pool anatomy in a macaque. **A**. Circles show averaged locations of the MN pools for each muscle in the transverse plane, the shaded areas are standard deviations along the dorsoventral and mediolateral directions calculated across all motoneuron locations in C5 through T1 segments. Grey matter outlines are shown for the caudal portion of the fifth cervical segment (C5C) and the caudal portion of the first thoracic segment (T1C, grey fill). The circle at the axis’s origin represents the central canal. **B**. 3D view of the individual motoneuron locations (colored circles) and grey matter outlines for spinal segments C5 - T1. Central canal locations are shown as open circles. Motoneuron pool colors are the same as in A. Insert on the right shows the rostrocaudal distributions of the motoneuron locations with black circles indicating averages.

To analyze the anatomical organization of MN pool locations, first the centers of MN pools were calculated by averaging the coordinates of all MNs comprising them. Then Euclidean distances were then calculated between each MN pool center. Unfortunately, the rostrocaudal distribution in our sample of MN pools was limited, with most centers of MN pools co-localizing between T8 and C1 (Fig. 1). Therefore, Euclidian distances were calculated not only in 3D, but also in 2D, with the rostrocaudal distribution excluded to control for the potential noise it introduced in our analysis.

### Musculoskeletal Models of Upper Extremity

We used open-source simulation software, OpenSim (version 4.1, Stanford University, Stanford, CA, USA), to model the musculoskeletal anatomy of both the macaque and human right upper limb. The macaque musculoskeletal model developed by Chan and Moran ^29^ was modified to add forearm muscles innervated by the MN pools included in the model described above. The following eight forelimb muscles have been added or modified based on anatomical data (Berringer, 1968): *adductor pollicis, dorsal interossei, extensor digitorum, extensor pollicis longus, flexor digitorum profundus, flexor digitorum superficialis, lumbricals*, and *ventral interossei* (Fig. 2A). Muscles with multiple heads and/or points of insertion were modeled as separate musculotendon actuators. For example, *biceps brachii* was modeled as two muscles, *biceps long head* and *biceps short head*. Similarly, the *extensor digitorum* was modeled as 4 muscles, sharing a similar origin, but inserting on each corresponding distal phalange of digits 2-5 (index through pinkie). Abbreviations of muscle names used in this study can be found in Table 1. To simulate more accurately the length changes of the additional muscles in different postures and during movement, we extended the model to include the carpometacarpal (CMC), metacarpophalangeal (MCP), and interphalangeal (IP) joints of the digits, increasing the total number of degrees of freedom (DOFs) in the macaque model to 27. The carpometacarpal joint of the thumb and the metacarpophalangeal joints of digits 2-4 were modeled with 2 DOFs, representing flexion/extension and abduction/adduction around the x- and z-axes, respectively. Interphalangeal joints were modeled as a single DOF representing flexion/extension about the x-axes.

**Figure 2.**
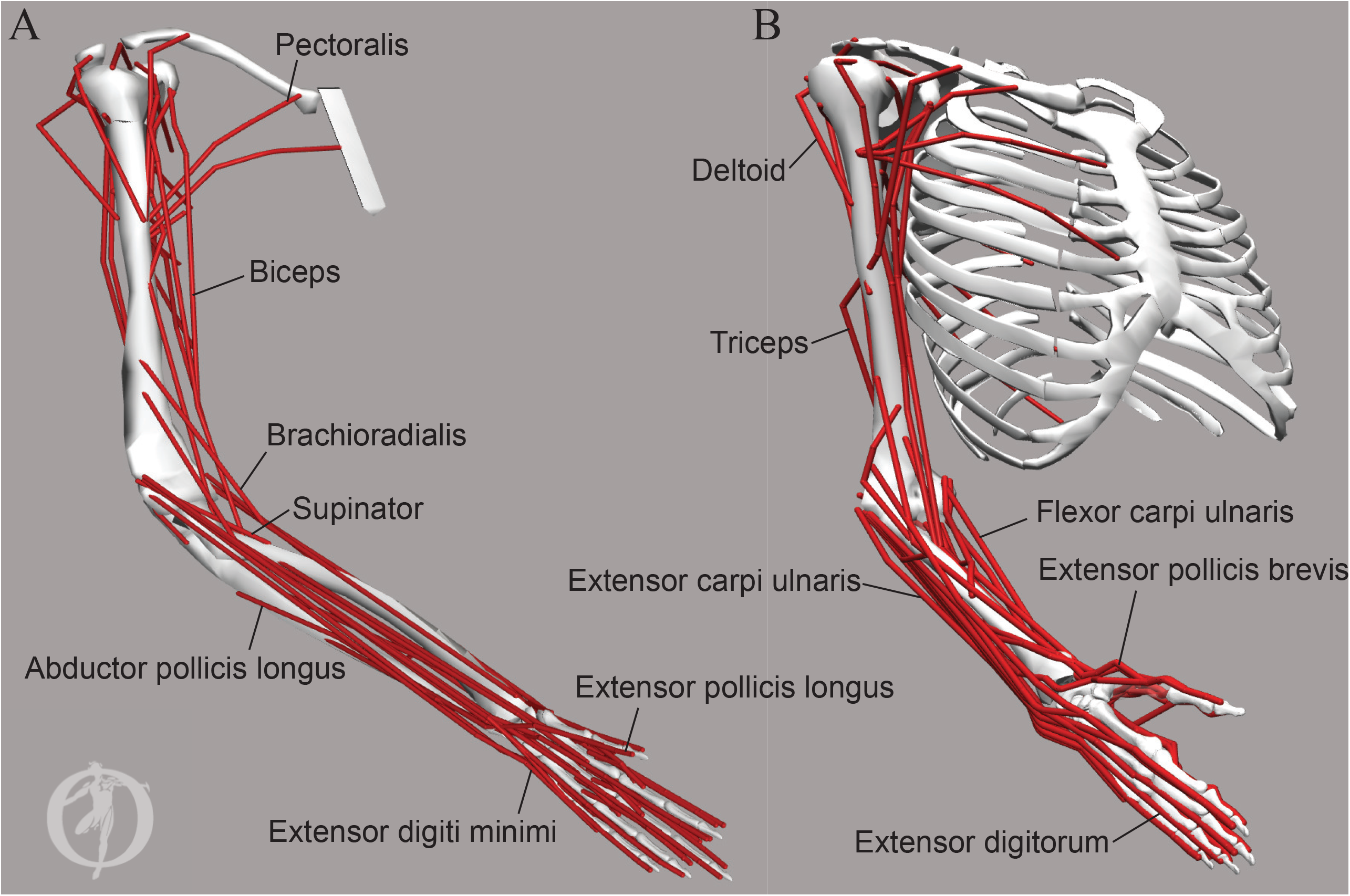
Musculoskeletal models of a macaque (A) and human (B) upper limbs. Red lines illustrate the origin, insertion, and wrapping geometry of the musculotendinous actuators representing the anatomical arrangement of individual muscles or muscle compartments. For example, two heads of biceps are modelled as two actuators with origin locations on different bones and a common insertion location. All actuators are shown, but not all are labeled. The bones are shown for illustration purposes only, the inertial geometries of limb segments to which the actuators attach are not shown.

**Table 1.**
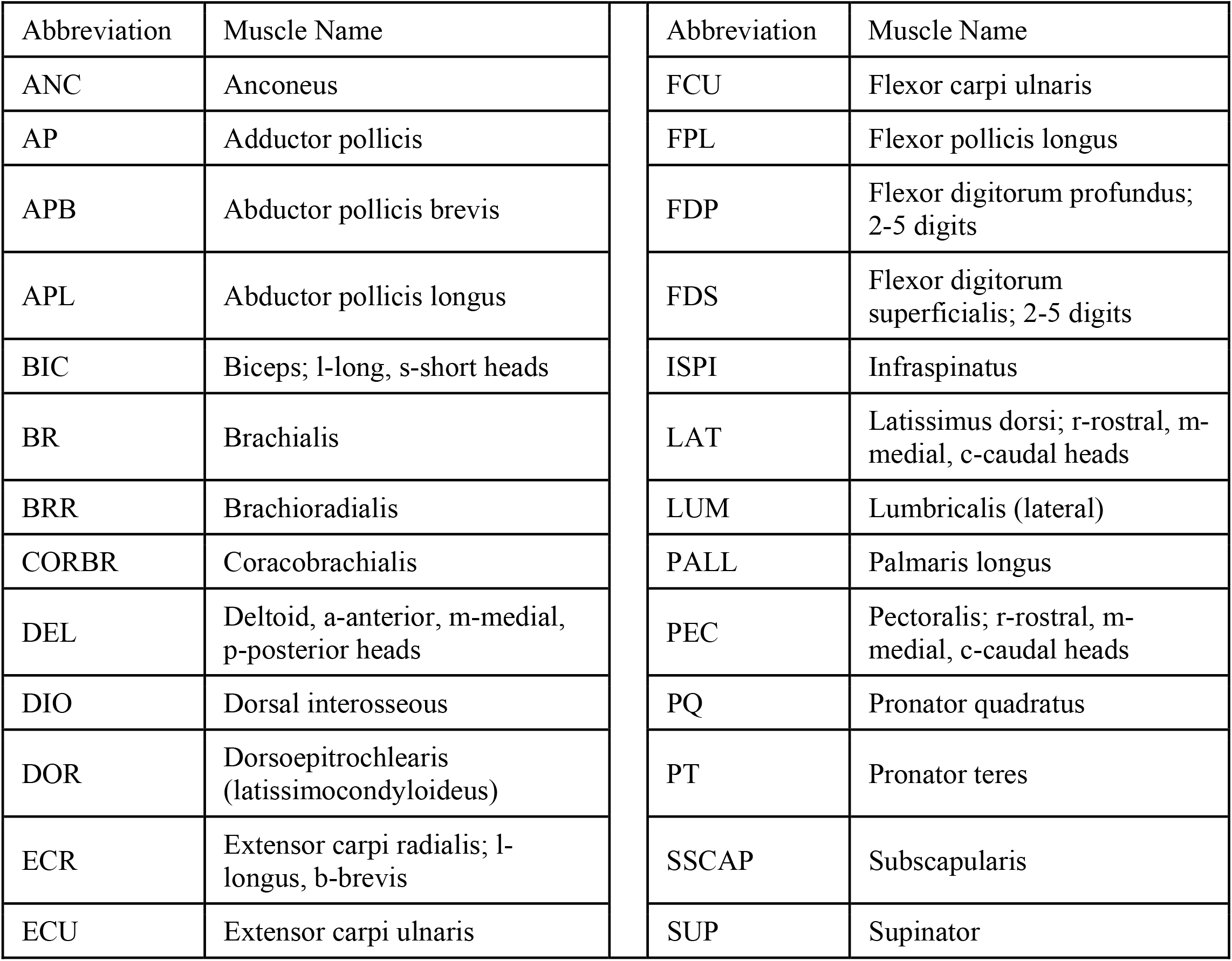
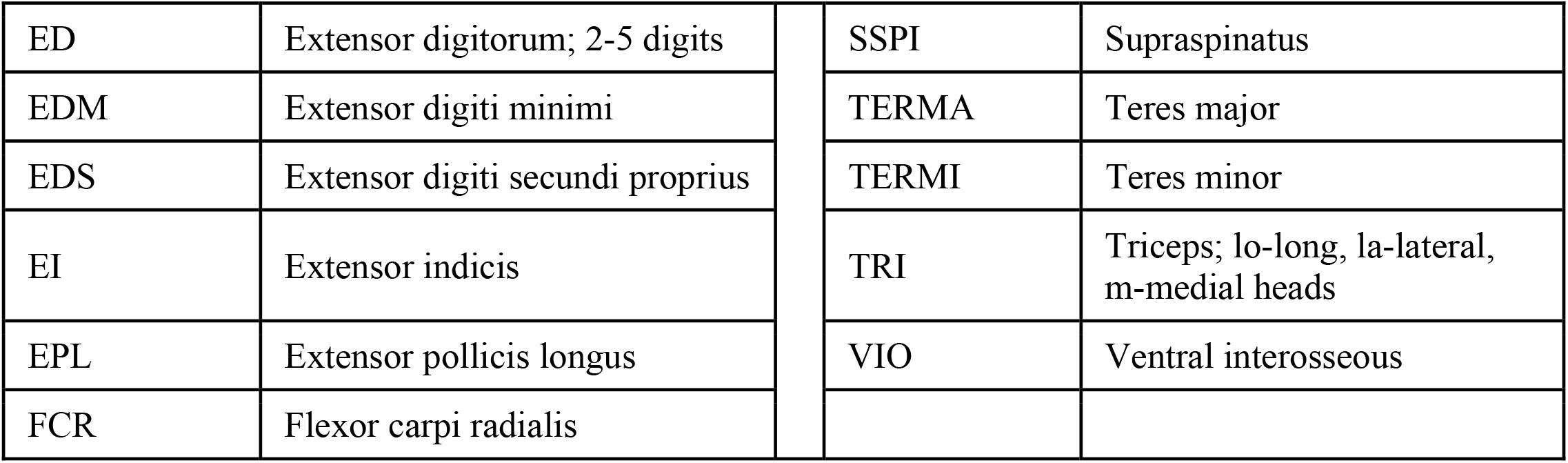
Muscle abbreviations used in this study.

A detailed model of the human arm was used to assess musculoskeletal differences between macaque and human musculoskeletal anatomy. The original model developed by Saul, et al. (2015) was previously expanded to include 23 DOFs and 52 musculotendinous actuators representing all major joints and 33 major muscles of the human arm (Gritsenko, et al. 2016) (Fig. 2B). The moment arms and maximal muscle forces were validated against published measurements ^30^. Only the DOFs spanned by muscles innervated by the MN pools modeled above were included in the comparative analysis of muscle lengths. These DOFs were elbow flexion/extension, hand pronation/supination, wrist joint flexion/extension and abduction/adduction, CMC joint flexion/extension and abduction/adduction, MCP joints 1-5 flexion/extension, and all IP joints 1-5 flexion/extension.

To analyze the functional organization of the musculoskeletal systems, musculotendon lengths were calculated for each forelimb posture across the physiological range of motion within the model. Then a correlation matrix of muscle lengths across all postures was then computed as in Gritsenko et al. ^16^. Positive correlations indicate a synergistic action of muscles, while negative correlations indicate antagonistic action. The correlations between muscle pairs whose MN pools were located closer vs. further apart were compared as described in the Statistical Analysis section.

### Statistical Analysis

The distances between MN pool centers were regressed against the correlation coefficients (R) between the lengths changes of muscles innervated by the corresponding MN pools. Nearby MN pools are more likely to share the descending inputs and afferent feedback, which makes them more likely to innervate synergistic muscles. Therefore, short distances that fell below the median of the distance distribution were analyzed separately from long distances that fell above the median. Thus, four regressions were done using 1) short 3D distances; 2) long 3D distances; 3) short 2D distances in the transverse plane; and 4) short 2D distances in the transverse plane. Familywise error for conducting the four tests was addressed using Bonferroni adjustment ^31^, alpha was set to 0.0125.

## Results

### Motoneuron clustering in the macaque spinal cord

The MN pools in the cervical enlargement of macaques were distributed along multiple segments (Fig. 1) similar to those in the lumbar enlargement of cats ^23^ and humans ^19,32^. MNs innervating proximal muscles in the macaque, such as the *biceps* and *triceps*, together with *flexor carpi radialis* and *extensor carpi radialis* were located in the rostral part of the cervical enlargement, while the rest of the MN pools were located in the caudal cervical and thoracic segments (Fig. 1B). In humans, the rostro-caudal distribution of MN pools has been estimated from the distribution of ventral roots across spinal segments and the muscles they innervate (Kendall and McCreary, 1983). The rostrocaudal distributions of human and macaque MN pools overlapped for all muscles (Fig. 3A). Although the MN pools innervating digit and some wrist muscles appeared to be more rostrally located and span more segments in humans compared to macaques. This may be due to the fewer number of spinal segments in humans compared to macaques ^33^, which may allow for more separation between MN pools in the rostrocaudal dimension in macaques. The distribution of MN pools in the transverse plane in macaque and human spinal cords also overlapped (Fig. 3B). In humans, the distribution of the MN pools in the transverse plane was estimated using postmortem staining by cresyl violet, as summarized in ^19^. The human MN pools in the lateral division of the ventral grey horn overlapped with those in the macaque (Fig. 3B). Although it was not possible to identify which muscles the human MNs innervated, the overlap with the macaque data suggests that the human MN pool labeled as 5 may innervate biceps and the human MN pool labeled as 7 may innervate triceps (Fig. 3B). Overall, these data show that there is a notable overlap between MN pools of humans and macaques.

**Figure 3.**
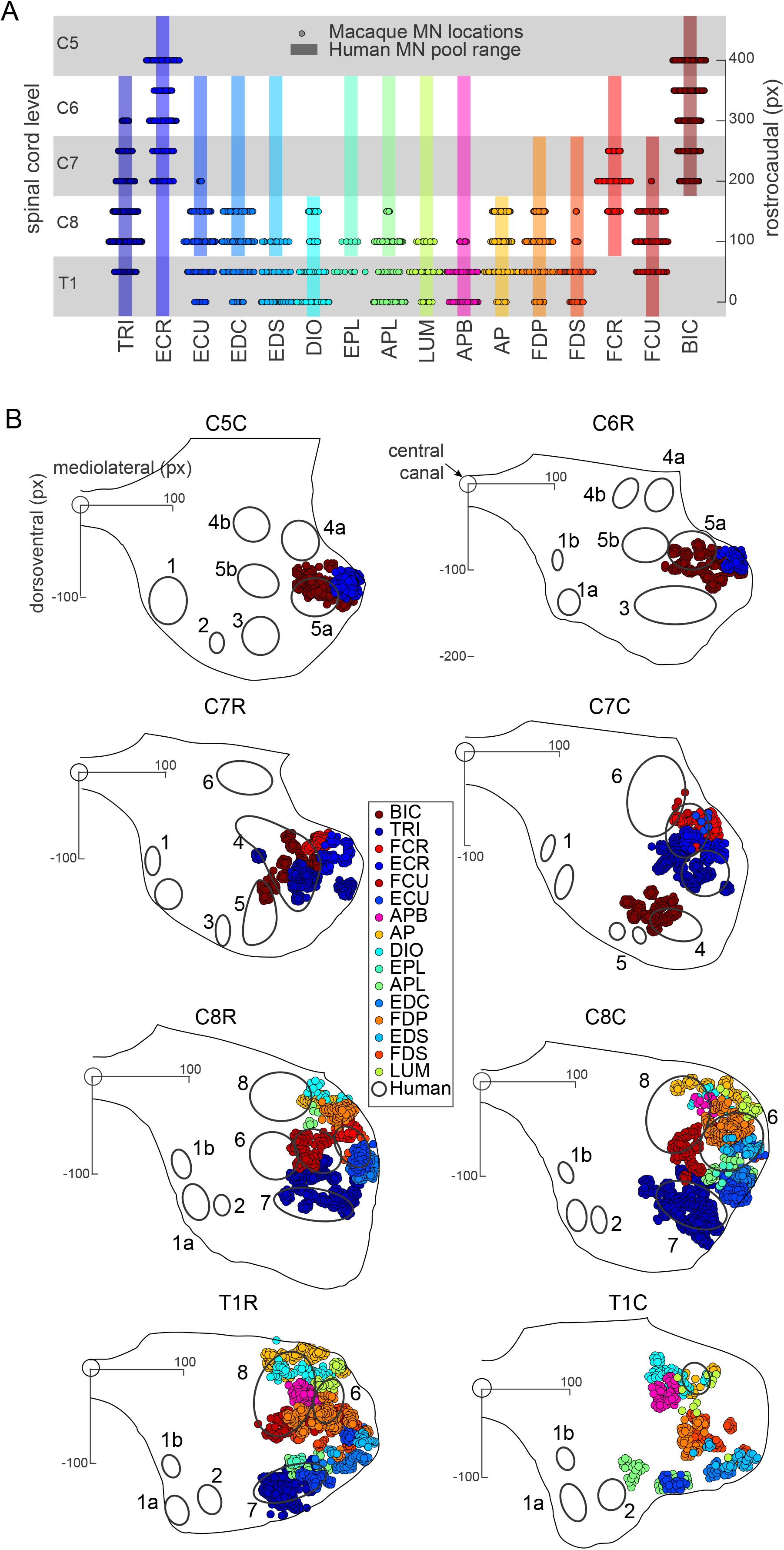
Comparison of MN locations between the macaque and human spinal cords. **A**. Circles show the distribution of MNs along the rostrocaudal direction across spinal segments from the macaque model in Fig 1. Bars show the distribution of corresponding MN pools from a human anatomy textbook ^32^. **B**. Circles show MN locations in the transverse plane per spinal segment from the macaque model in Fig 1. Grey matter outlines aligned on the central canal at the origin of the axes are from the macaque model. Black open ovals show the distributions of select MN pools adapted from human tracing studies summarized in ^19^. The human distributions are numbered as in the Routal and Pal (1999).

To quantify the anatomical organization of spinal MN pools we calculated the Euclidian distances between the centers of MN pools. The MN pools innervating *biceps* and *extensor carpi radialis* were the furthest apart from the rest of the MN pools, while being a relatively short distance from each other (Fig. 4A, dark red cluster in the top left corner). Moreover, several other MN pools innervating synergistic muscles were co-localized, such as the MN pool of *flexor digitorum superficialis* was a relatively short distance apart from that of *flexor digitorum profundus*. The MN pools innervating two long thumb muscles (*extensor* and *abductor pollicis longus*) were co-localized. The MN pools innervating two other thumb muscles (*adductor pollicis* and *abductor pollicis brevis*) were also co-localized with those innervating intrinsic hand muscles (*dorsal interosseous* and *lumbricalis*) (Fig. 4A, dark red cluster in the bottom right corner). The close anatomical arrangement of MN pools remained the same when the rostrocaudal distribution was excluded from the distance calculations (Fig. 4B). This anatomical arrangement suggests that the MN pools innervating agonistic muscles that perform a single function, such as spreading fingers apart in the case of the last cluster, are located closer together than the MN pools innervating antagonistic muscles that perform distinct functions. We have previously conducted a rigorous analysis of the agonistic and antagonistic relationships between muscles using a model of musculoskeletal anatomy of the human arm ^16^. Here we have included a musculoskeletal model of the macaque upper limb and compared the functional relationships between muscles to the anatomical relationships between MN pools that innervate those muscles.

**Figure 4.**
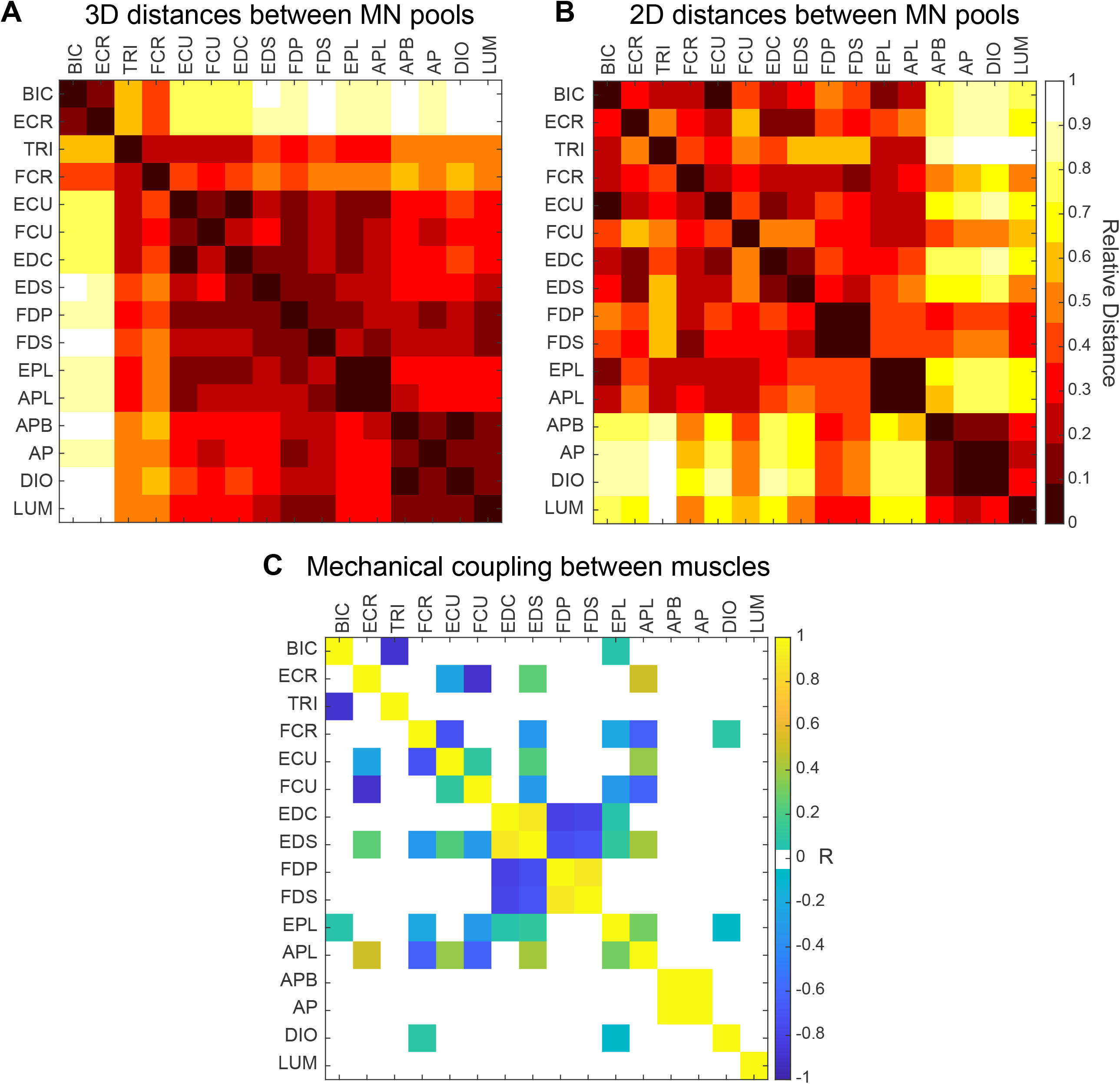
Anatomical relationships of MN pools and the functional relationships between muscles they innervate. Correlation matrix of the relative distances between the centers of MN pools in all dimensions (**A**) and only in the transverse plane with rostrocaudal distribution excluded (**B**) calculated using the model shown in Fig. 1. Distances were calculated between the centers of MN pools for each pair of MN pools relative to the maximum distance across all pairs. Black indicates short distances, white indicate long distances. **C**. Matrix of Pearson correlation coefficients (R) between muscle length changes evaluated across the full range of motion of the macaque forelimb. Yellow indicates agonistic actions; blue indicates antagonistic actions.

To evaluate the functional relationship between muscles we correlated changes of muscle lengths across the whole range of possible upper limb postures ^16^. The logic here is that muscle length changes that are positively correlated indicate agonistic action, while muscle length changes that are negatively correlated indicate antagonistic action. For example, during wrist extension *extensor carpi ulnaris* and *extensor digitorum communis* would shorten and their lengths would be positively correlated indicating their agonistic action, while *flexor digitorum profundus* and *flexor digitorum superficialis* would lengthen and their lengths would be negatively correlated with that of the former two muscles indicating their antagonistic action. Indeed, all muscles but *dorsal interosseous* and *lumbricalis* had strong agonistic or antagonistic relationships with at least one other muscle based on the length comparisons using the macaque musculoskeletal model (Fig. 4C). This shows that there are strong functional relationships between most muscles of the upper limb that are innervated by the MN pools included in the spinal cord model.

Moreover, the functional relationships between all major muscles of the upper limb of the macaque were very similar to those observed in human arm (Fig. 5). This shows that the functional relationships are preserved across species with similar anatomy.

**Figure 5.**
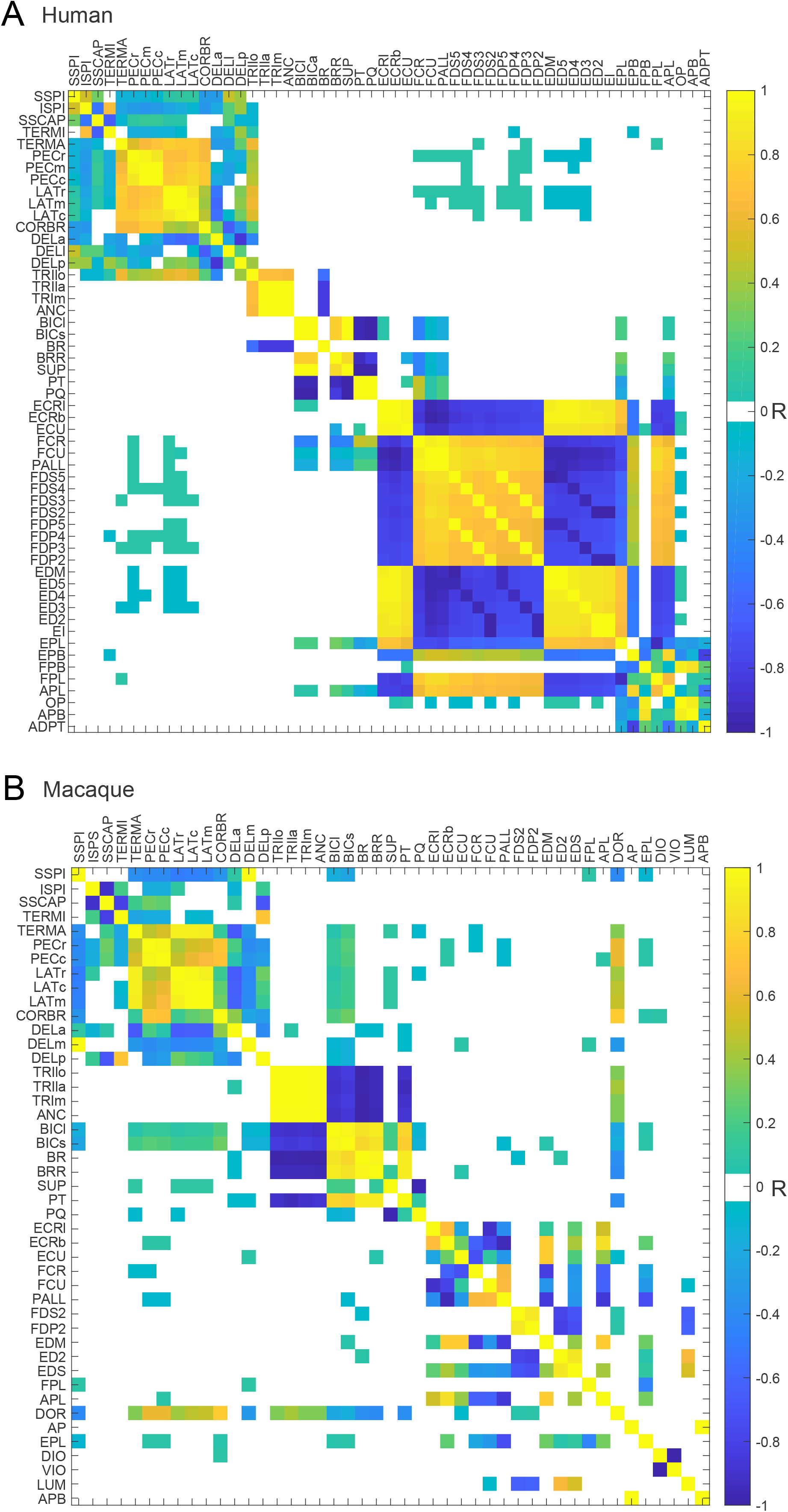
The functional relationships between upper limb muscles of human and macaque. Matrices of Pearson correlation coefficients (R) between muscle length changes evaluated across the full range of motion of the human arm **(A)** and macaque forelimb **(B)** shown in Fig. 2. Yellow indicates agonistic actions; blue indicates antagonistic actions as in Fig. 4C.

Next, we compared the functional relationship between muscles with the spatial relationship between MN pools innervating these muscles. We examined separately the 3D Euclidian distances between MN pool centers that fell below and above the median of the distribution (Fig. 6A). There was no significant linear regression between the relative locations of nearby MN pools and the functional relationships of the corresponding muscles (Fig. 6B). However, when the Euclidian distances were calculated only in the transverse plane, i.e., excluding the rostrocaudal distribution (Fig. 6C), a significant linear regression was revealed between the relative locations of nearby MN pools and the functional relationships of the corresponding muscles (Fig. 6D). No significant regressions were found for MN pools that were located further apart from each other (3D locations: R^2^ = 0.127, *p* = 0.16; 2D locations: R^2^ = 0.071, *p* = 0.57).

**Figure 6.**
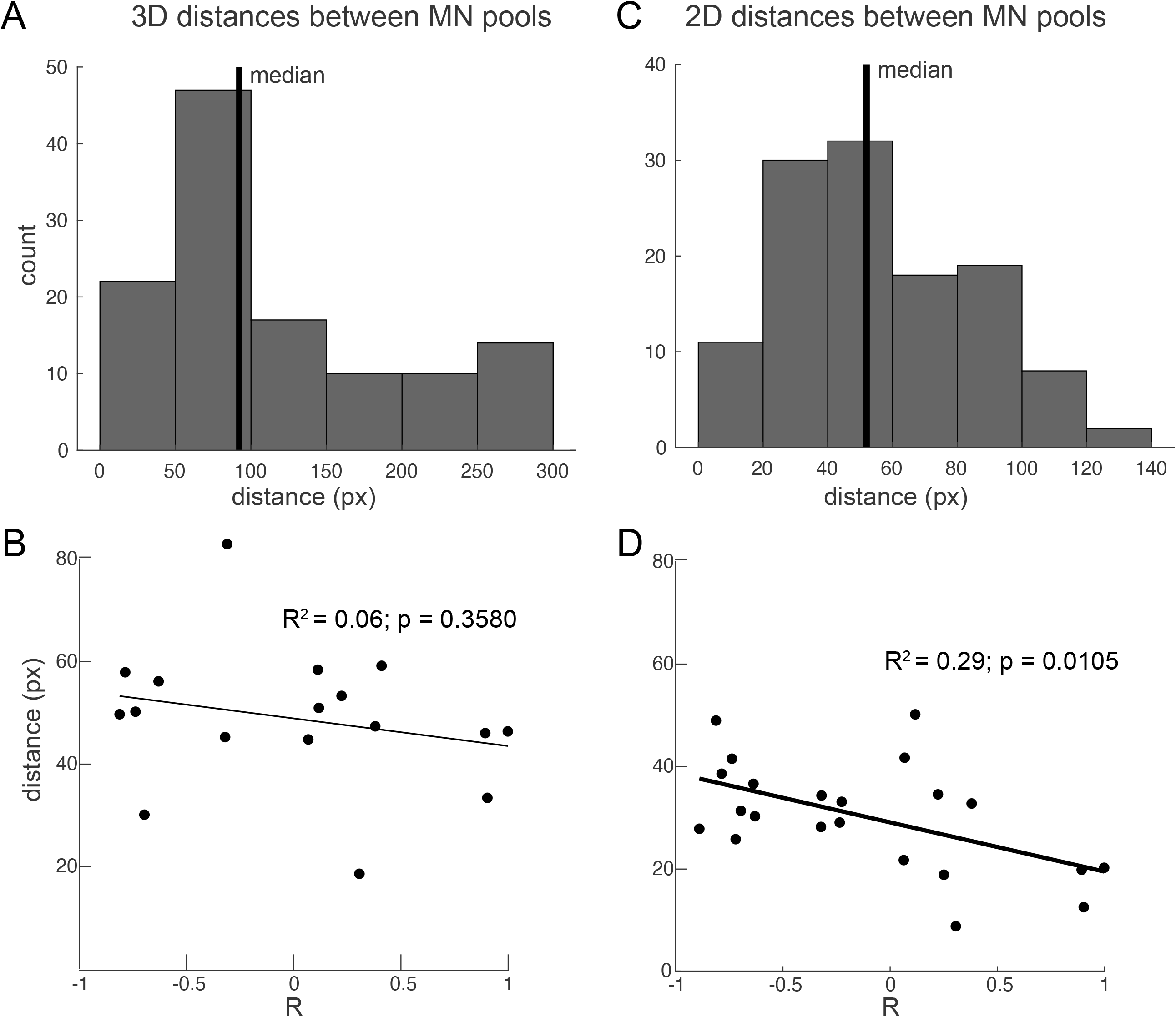
Comparative analysis of spatial MN pool anatomy and functional relationships between muscles they innervate. **A**. Histogram of 3D distances between MN pools. Vertical black line shows the median, only distances below the median were used for the regression in B. **B**. Short 3D distances between MN pools as a function of Pearson correlation coefficients (R) between lengths of muscles those MN pools they innervate. The regression line is not significant as shown by the inserted statistics. **C**. Histogram of 2D distances in the transverse plane between MN pools. Vertical black line shows the median, only distances below the median were used for the regression in D. **D**. Short 2D distances between MN pools as a function of Pearson correlation coefficients (R) between lengths of muscles those MN pools they innervate. The regression line is significant as shown by the inserted statistics.

This shows that the spatial distribution of the MN pools in the transverse plane mimics the functional relationships between the muscles they innervate.

## Discussion

Here, we have developed the first spatial model of cervico-thoracic MN distributions in the macaque. Using this model, we have shown that the MN distributions in the macaque spinal cord are broadly similar to those in human spinal cord (Fig. 3). Using the musculoskeletal models of the macaque and human upper limbs we have also shown that the muscle anatomy of both species serves similar agonistic and antagonistic actions (Fig. 5). We then demonstrated quantitatively that the spatial distribution of MN pools co-localized in the transverse plane mimics the functional relationships between the muscles these MNs innervate (Fig. 4, 6). This suggests that the spatial organization of the MN pools embeds the functional musculoskeletal anatomy evolved to support the large repertoire of human arm movements ^16^.

The functional organization of the spinal cord is modular ^34^. This conceptual understanding emerged from early studies in spinalized animals showing that microstimulation inside the spinal cord grey matter produces contractions of multiple muscles, which cause converging forces toward a subset of postures ^35^. The idea then emerged that muscles can be controlled in groups with fewer control signals of varying spatial and temporal patterns that can create a large repertoire of movements ^36^. There is also a parallel line of investigation of the spinal control of locomotion that reveals the modularity of spinal control circuitry including the central pattern generator (CPG) ^37^. Evidence supports the importance of CPG in human sensorimotor control ^4,38,39^. There is evidence that the different spatiotemporal dynamics of the CPG and its interaction with the body through afferent feedback can generate different behaviors. This can be seen in neuromechanical studies with lampreys where varying the spatiotemporal patterns of tonic excitation to CPG and its propagation along the spinal segments can produce swimming in either forward or backward direction ^40^ and change the speed, direction and type of gait ^41,42^. Some of the spatiotemporal dynamics of the CPG during gait has been shown to be reflected in the anatomical distribution of the MN pools innervating hindlimb muscles of the cat ^23^, supporting the idea of the somatotopic “map” of a hindlimb ^43^. Our study provides supporting evidence for the idea of the somatotopic “map” of a forelimb. Our results show that the MN pools innervating muscles that perform synergistic action are co-localized in the spinal cord. Nearby MN pools are also likely to share the proprioceptive feedback from the muscles they innervate as has been shown for cat knee extensor muscles and their afferents ^44^. Similarly, nearby MN pools are likely to receive common descending signals from propriospinal interneurons ^45^ that can explain observations of the convergent force fields that are thought to serve as building blocks for more complex motor control ^35,46^. Furthermore, nearby MN pools are likely to receive common descending signals from the somatotopically organized motor cortex that can explain observations of the complex movements in different areas of workspace produced as a result of the stimulation of the primary and premotor cortex ^47^. Overall, our results suggest that the spatial organization of MN pools may play an important role in neuromechanical tuning and simplify the sensorimotor control of the upper limb.

Recent work has shown that it is possible to restore trunk and leg motor functions using biomimetic epidural electrical stimulation of the dorsal roots below the lesion caused by the spinal cord injury ^7^. Our results suggest that a similar approach may work for restoring complex movements of the arm after spinal cord injury. A proof of concept study in macaques have shown that it is possible to evoke selective muscle responses by stimulating cervical dorsal roots ^48^. Notably, more rostral stimulation in this study evoked responses in *biceps* and *extensor carpi radialis*, while more caudal stimulation evoked responses in *flexor digitorum superficialis, extensor digitorum communis*, and *abductor policis brevis*. The rostrocaudal distribution of MN pools innervating these muscles is similar to that shown by our model (Fig. 3A). These observations support the validity of our model and indicate potential future uses of it for designing next generation neuroprosthetics.

## Acknowledgements

We would like to thank Ariel Thomas and Brian Tomblin for digitizing images from the paper and Mark Kozy for scripting help. VG & SY were supported by NIGMS grants P20GM109098 and P30GM103503.

